# SYRINGAE: A web-based application for *Pseudomonas syringae* isolate characterization

**DOI:** 10.1101/2022.11.04.515192

**Authors:** Chad Fautt, Estelle Couradeau, Kevin L. Hockett

## Abstract

The *Pseudomonas syringae* species complex (PSSC) is a diverse group of plant pathogens with a collective host range encompassing almost every food crop grown throughout the world. As a threat to global food security, rapid detection and characterization of epidemic and emerging pathogenic lineages is essential. However, phylogenetic identification and prediction of virulence is often complicated by an unclarified taxonomy and the diversity of virulence factor repertoires carried by PSSC isolates. To address these issues, we have built SYRINGAE (www.syringae.org), a web-based phylogenetic placement and functional inference pipeline for PSSC. SYRINGAE contains a comprehensive phylogeny of 2,161 quality-checked genome assemblies annotated with 120 virulence genes. From this dataset, naïve Baye classification models trained from life identification numbers (LINs) and common marker gene sequences can be used for accurate identification of isolates. SYRINGAE efficiently articulates taxonomical and functional data generated over the last several decades on PSSC and constitutes a unique tool tailored towards the rapid characterization of PSSC emerging strains of concern.

## Background & Summary

The *Pseudomonas syringae* species complex (PSSC) has been co-evolving with plants since before the emergence of angiosperms (Xin, Kvitko and He, 2018), and has diversified into one of the most economically important groups of plant pathogens in the world, with a collective host range spanning almost every major food crop grown today (Baltrus, McCann and Guttman, 2017). Critically, while there are many pathogens within PSSC, there is also a wide range of virulence exhibited throughout the species complex, including non-pathogenic plant epiphytes and clades associated with the water cycle (Morris *et al.*, 2007, 2008). The ability to discriminate between lineages within the PSSC and rapidly predict potential pathogenicity of novel lineages is crucial for preventing epidemic outbreaks (Cunty *et al.*, 2015), detecting emerging pathogenic strains (Zhao *et al.*, 2022), and untangling correlations between virulence factors carried by a pathogen, its host range, and its virulence (Preston, 2000). Although the efforts to catalog PSSC diversity and to understand the molecular determinants of virulence have yielded great insights into their ecology and behavior (Morris, Monteil and Berge, 2013), currently there is no efficient way to leverage these insights to efficiently predict the identity and pathogenicity of newly discovered PSSC strains. This is especially true for those researchers or labs that do not specialize on PSSC.

A major barrier to the characterization of PSSC strains is the inconclusive or inaccurate taxonomic identities of published genomes. By one estimate, 42% of all published PSSC genomes are misclassified at the species level, based on analysis of phylogenetic relationships described by average nucleotide identity (ANI) and multi-locus sequence analysis (MLSA) (Gomila *et al.*, 2017). As genomes deposited in databases such as GenBank often serve as reference sequences for identification of isolates found on or near diseased plants, the high rate of misclassification has a direct, negative impact on our ability to efficiently recognize pathogenic lineages. The designation of 13 phylogroups based on MLST has clarified phylogenetic relationships within PSSC (Berge *et al.*, 2014), however most published genomes aren’t ascribed to a phylogroup in public databases and thus their use in classification is limited. While Berge et al. have addressed this shortcoming by providing a reference database of phylogroup type strains allowing classification based on the CTS gene, there has since been no broader effort to make the classification process more efficient. Yet another approach to circumvent the inaccurate taxonomy at the species level while allowing for placement into clades below the species and phylogroup level is the clustering of genomes by ANI (Average Nucleotide Identity) (Vinatzer *et al.*, 2017). This approach assigns a life identification number (LIN) to each unique genome in a database, creating hierarchical clusters of genomes that largely recapitulate traditional phylogenetic clades described by the core genome, and allow for higher resolution than traditional PSSC taxonomy (Figure 1).

**Figure 1.**
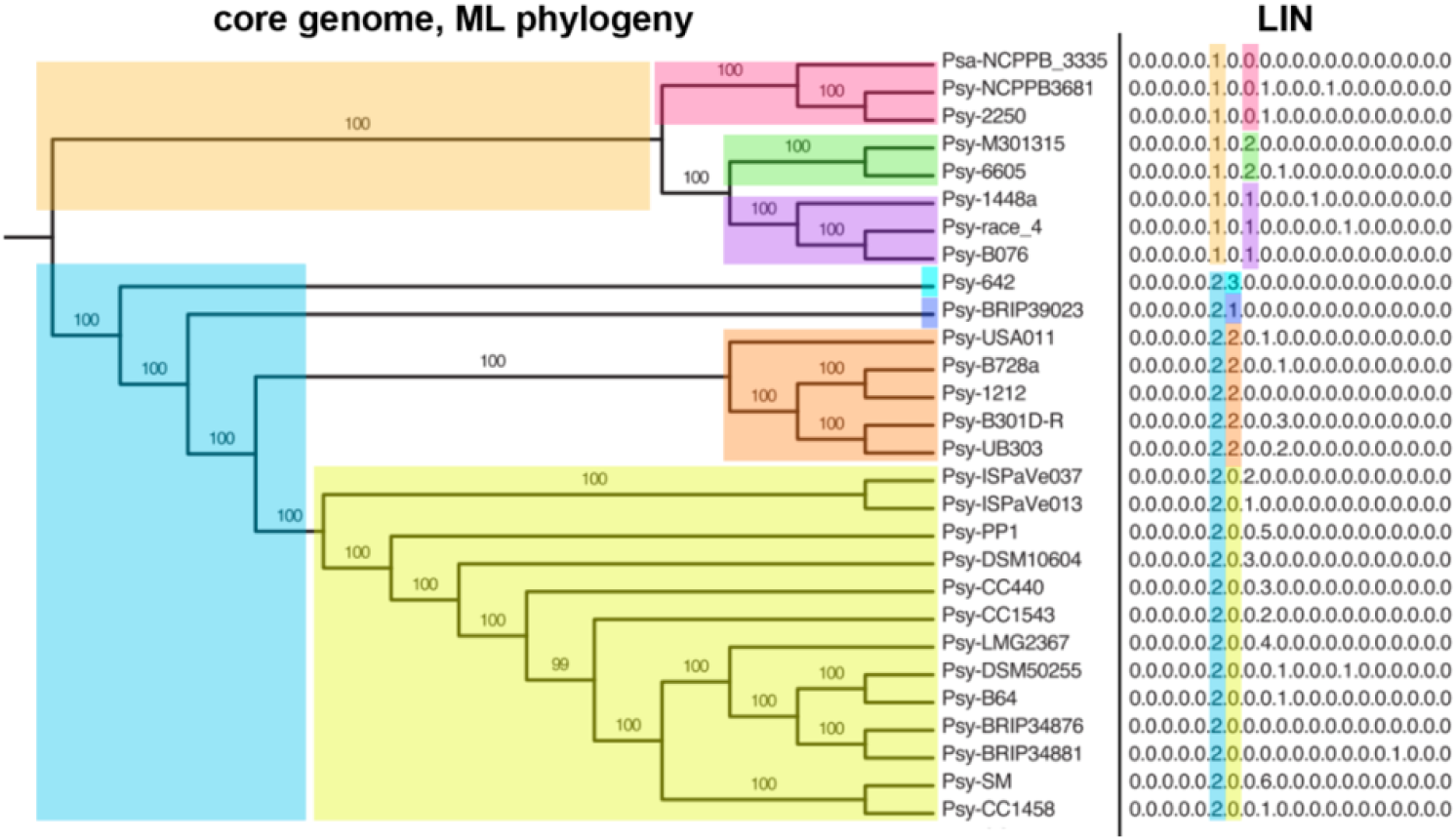
Comparison of clustering within PSSC that results from a maximum likelihood phylogenetic tree and LIN assigned based on ANI. Digits from left to right in each LIN correspond to inclusion of a strain in increasingly smaller clades within the phylogeny. Figure adapted from Vinatzer et al., 2017

A second barrier to characterization of new PSSC isolates is the functional diversity exhibited throughout the species complex. Specifically, host range and virulence can vary considerably among pathogenic strains belonging to the same pathovar, while strains belonging to different pathovars can nonetheless exhibit similar host ranges. These complex patterns stem, at least in part, from the formal definition of pathovar as ‘a strain or set of strains with the same or similar characteristics, differentiated at infrasubspecific level from other strains of the same species or subspecies on the basis of distinctive pathogenicity to one or more plant hosts.’ which allows for broad interpretation. As such, some pathovars, such as pv. avii, have been described by their ability to cause disease on a single host (10.1023/a:1024786201793), some as having different host ranges among a small defined group of hosts (P. savastanoi pvs. savastanoi, nerii, fraxini, mandevillae and retacarpa, https://doi.org/10.1094/PHYTO-11-20-0526-R), while it has also been argued that pathogens sharing a wide common host range, regardless of a shared pathogenic potential for any single host, should also be considered as belonging to a single pathovar (Young, 2008). Given the inconsistent criteria for delineating between pathovars, and recent evidence that host ranges in PSSC strains overlap with no discernable modularity (Morris et al., 2019), some groups have called into question the validity of pathovar designations for epidemiological and disease management purposes (Lamichhane et al, 2014). Further, properly assigning a given isolate to an appropriate pathovar requires performing host range tests that are prohibitively laborious to many labs. An alternative phylogenomic approach to predicting pathogenic potential would be beneficial, as others have demonstrated that comparative genomics can discriminate between strains known to have different host ranges (Moreno-Pérez et al., 2020) and correctly identify strains capable of infecting a given host (Almeida et al., 2022). In both of the above cases, known virulence factors, particularly those associated with the type III secretion system (T3SS), were highly correlated with known virulence patterns. Assuming T3SS effector proteins are conserved at some phylogenetic level, these results indicate that a phylogenomic signal may be present in PSSC that would be useful for assessing pathogenic potential without laborious experimental assays.

In a recent contribution we showed that we could accurately predict the presence of 77 type III effector subfamilies in PSSC isolates with an accuracy of 79-94% (REF), using only single amplicon sequence data. Here, we introduce SYRINGAE, a web application for phylogenetic classification and prediction of carried virulence factors for unknown PSSC isolates using user-submitted amplicon data generated from common PCR primer sets. To build the underlying databases of SYRINGAE, 2,161 quality-checked genomes downloaded from GenBank were screened for known virulence factors associated with the T3SS, type 3 effectors (T3E), and the Woody Host or Pseudomonas (WHOP) region associated with knot formation in woody hosts (Caballo-Ponce *et al.*, 2017). Pairwise ANI values were calculated for all genomes, and a maximum-likelihood phylogeny was constructed from the core genome of PSSC. Using the LIN clustering algorithm as a strict hierarchical classification scheme, classification models were trained and made available on the SYRINGAE website. We believe this resource will be particularly useful for researchers who are new to working with PSSC isolates and do not already specialize on the *P. syringae* group as a focal system.

## Methods

### reference PSSC genomes

All genome assemblies classified as *‘Pseudomonas syringae* group’ were downloaded from the GenBank via NCBI in November 2021. Genomes were checked for completeness and assembly quality with BUSCO v5.3.1 using the pseudomonadales_odb10 lineage database, and genomes with a BUSCO score ≥ 99 were kept for further processing (Fig 2a). A CSV file summarizing each genome was created (and made searchable on the SYRINGAE website) and available at xxx. Data included in this file are NCBI-submitted taxonomic data, type strain designations, phylogroups as assigned in this study, LIN clusters assigned for classification purposes, presence/absence of key virulence factors, and metadata found in each genome’s Biosample record.

**Figure 2.**
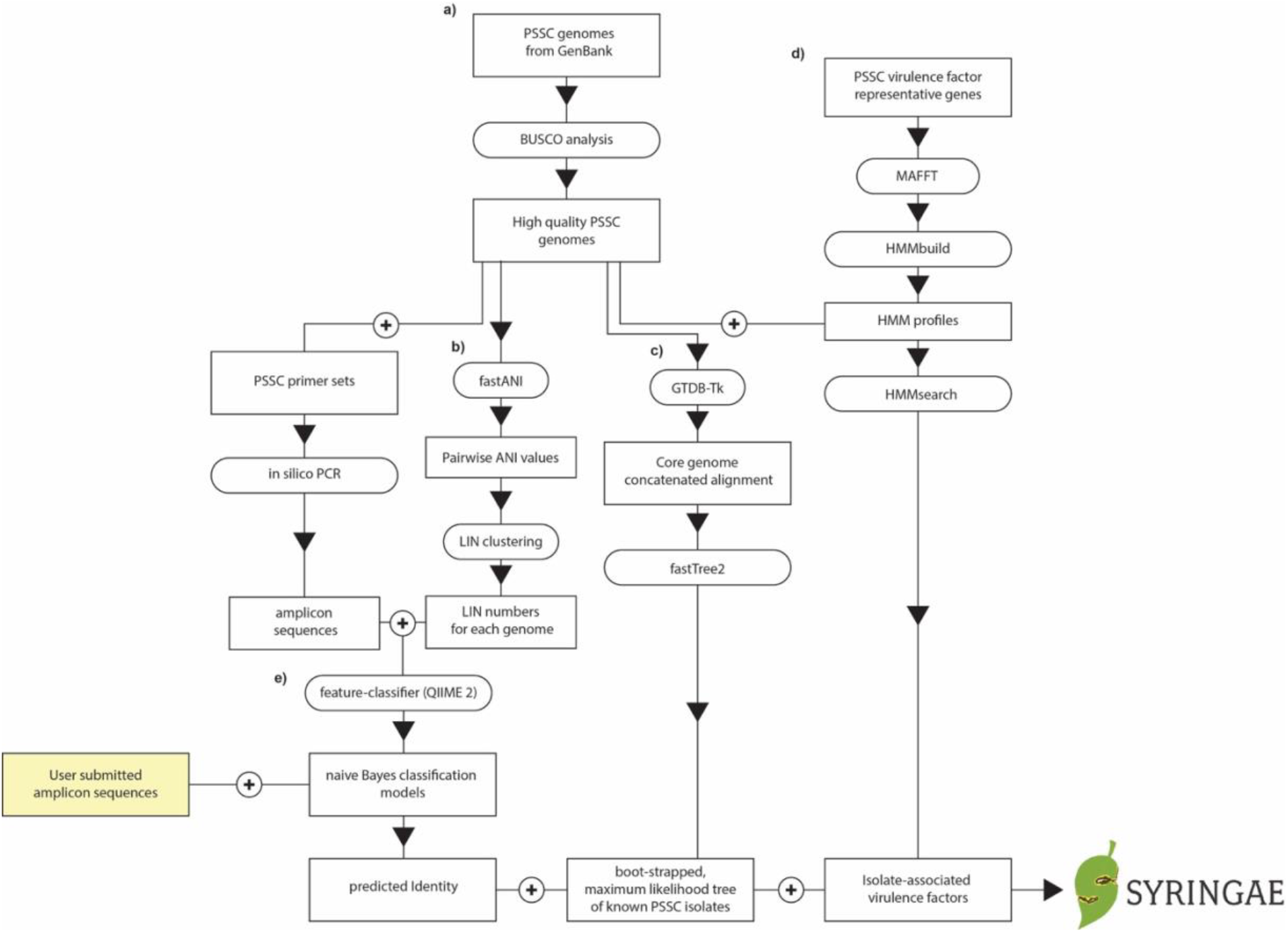
Schematic of bioinformatic pipeline used for SYRINGAE.

### Assigning phylogroups to genomes

Phylogroup assignment of each genome was based on ANI shared with previously classified reference strains representing Phylogroups 1a,1b,2a,2b,2c,2d,3,4,5,6,7,9,10,11, and 13 (Berge *et al.*, 2014) (supplementary table 1). Reference strains for phylogroups 8 and 12 were not found among the 2,161 genomes characterized by SYRINGAE, either because they were not represented in the GenBank database or did not make it past the BUSCO quality check described above.

A genome was assigned to a given phylogroup if it was the most closely related to the reference strain for that phylogroup, based on ANI. To minimize inaccurate phylogroup assignments, any genome sharing less than 95% ANI to any reference strain was left unassigned.

### Assigning LIN clusters to genomes

A significant barrier to PSSC classification is unreliable and inconsistent taxonomic assignments. As such, SYRINGAE utilizes hierarchical clustering based on ANI values as an alternative to the Linnean taxonomy files typically used for Bayesian classification. Pairwise ANI between all genomes was calculated using fastANI v1.33 with default settings. Using the algorithm previously described (Vinatzer *et al.*, 2017), each genome was assigned to LIN cluster (fig 2b). To describe the algorithm briefly, a random genome was designated as belonging to group ‘0’ at every ANI bin (e.g. assuming ANI bins of 80, 90, and 95% would give a LIN number of ‘0.0.0’). Each subsequent randomly selected genome was assigned a LIN number based on the genome it has the highest ANI with among genomes already assigned a LIN number. If, for example, the second genome selected had an ANI of 92% with the first genome, its LIN number would be assigned as ‘0.0.1’, as it meets the threshold for belonging to the same group as the first genome at the 80 and 90% ANI levels but differs from the first genome at the 95% level, and so a new group ‘1’ is created for it. All genomes were sequentially assigned LIN numbers in this way. For SYINGAE, ANI bins at 1% increments between 80-99% were used. A drawback to using LIN clustering for classification is the LIN number assigned to a given genome is highly dependent on the order of genomes selected for clustering. Thus, results obtained from classification models built with a LIN ‘taxonomy’ can only be interpreted when used in conjunction with a database outlining LIN number the classifier was trained on. The SYRINGAE website overcomes this limitation by using a precomputed unique set of LIN numbers for accurate classification purposes and by conveying the results to the user in an interpretable format. To achieve that, SYRINGAE 1) cross-references the predicted cluster represented by a given LIN number with traditional taxonomy and 2) displays its position on a whole-genome phylogenetic tree.

### Building Phylogenetic tree

A concatenated gene alignment based on the core genome of PSSC was generated using GTDB-TK (Chaumeil *et al.*, 2020). From this alignment, FastTree2 (Price, Dehal and Arkin, 2010) with default settings was used to construct an approximately maximum-likelihood phylogenetic tree (fig 2c).

### Screening genomes for virulence factors of concern

Canonical T3SS genes from PSSC strains DC300 and B728A were used to seed a search for homologs with an 85% identity threshold using the Geneious prime 2019 software package (Dotmatics, 2022).The same method was used to find representative WHOP region gene sequences, using previously annotated genes in strain NCPPB 3335 (Caballo-Ponce *et al.*, 2017) as seeds for the homology search. T3E reference sequences were obtained from Psytec (Laflamme *et al.*, 2020). Reference sequences obtained for each gene were aligned with MAFFT (Katoh and Standley, 2013) using default settings, and alignments were used as input for a more robust identification of homologs in the entire set of 2,161 PSSC genomes using HMMER v3.3.2 (Eddy, 2020) (fig 2d). HMMER output files were manually inspected and based on the E-values associated with known virulence factors, matches with an E-value < 10^-20^ were considered to be statistically significant hits. In instances where two genes were identified as more than one virulence factor (a common occurrence among closely related T3E subfamilies), the identification with the lowest E-value was chosen as the official annotation.

### PSSC primer sets supported by SYRINGAE

Over the last two decades, several PCR primers have been developed, often as part of MLST schemes, for building evolutionary accurate phylogenies and aiding in classification of unknown isolates. More recently, there has been interest in utilizing single amplicon sequences for these purposes. To help in our decision of which primer sets to include in SYRINGAE, as well as to investigate which provide the most value for classification using a single amplicon, we conducted a short but thorough in-silico investigation of 16 commonly used primer sets (Fautt et al., submitted). Briefly, we assessed successful amplification in genomes representing the full diversity of the species complex as we currently know it, concordance between pairwise amplicon distance and whole genome ANI, resolution of naïve Bayes classifiers trained from amplicon data, as well as the potential for functional prediction based on the classification results. Given the results of that study, SYRINGAE supports the following primer sets: GyrB, GapA, CTS, RpoD, (Hwang *et al.*, 2005) and PGI (Yan *et al.*, 2008) (Table 1).

**Table 1.**
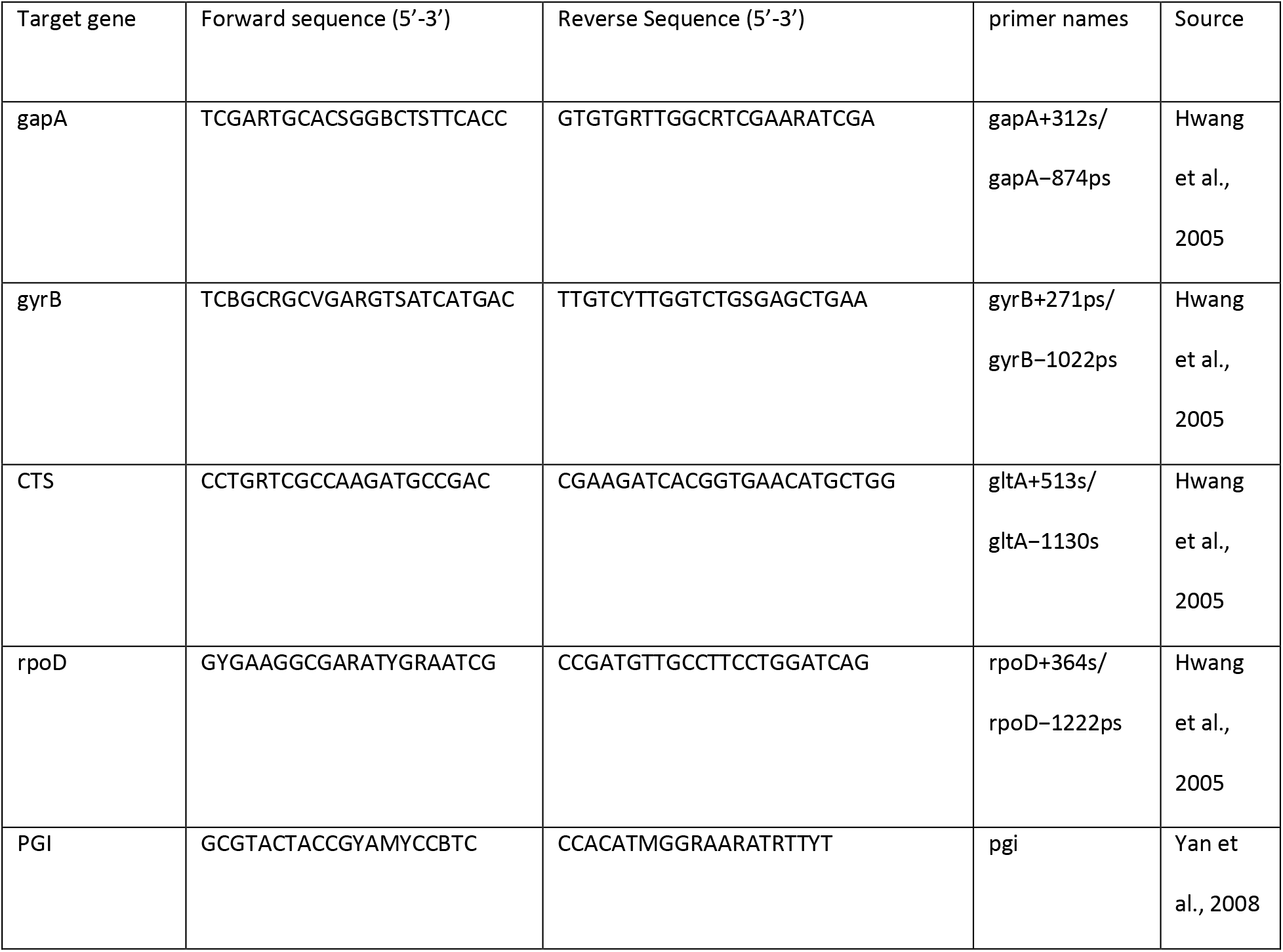
primer sets accepted by Syringae.org for isolate characterization.

### Training Naïve Bayes classification models

For each marker gene, a classification model was trained using the feature-classifier plugin for QIIME 2. Training naïve Bayes classifiers requires both a list of sequences, and an associated taxonomy file for each sequence (typically in the format ‘Order_Pseudomonadales; Family_ Pseudomonadaceae; Genus_Pseudomonas; Species_syringae;’). LIN numbers assigned to each genome were used to construct a hierarchical taxonomy, with ANI bins within each LIN number acting as taxa levels, and groups acting as individual taxa (e.g., a taxonomy format of ‘80%_0; 90%_0; 95%_1) (fig 2e).

### Technical Validation

Genome records included in SYRINGAE were validated for assembly quality using BUSCO, and all genomes scoring < 99 were removed from the dataset. Accuracy of the classification models and functional predictions were investigated and published separately (Fautt et al., submitted). Beyond the T3SS, T3E, and WHOP genes, which were annotated using HMM models built for this study, all gene annotations were taken directly from the NCBI Prokaryotic Genome Annotation Pipeline.

### Usage Note

#### Identify

The primary functionality on syringae.org is the rapid characterization of *Pseudomonas sp*. isolates from single amplicon sequences.

Input:

> The query sequence(s) are untrimmed amplicon sequences generated from primer sets targeting the genes gyrB, CTS, gapA, PGI, or rpoD (table 1), in FASTA or multiFASTA format.

Outputs:

1. A list of genomes predicted to be most closely related to the unknown isolate
2. Phylogenetic classification for each query sequence is also displayed as a phylogenetic tree rooted at the most recent common ancestor of all genomes in the Syringae database predicted to be closely related to the unknown isolate. By default, the tree only shows the most closely related genomes, but users can toggle the ANI threshold scale to lower values, to “zoom out” and display more distant relatives. This extra functionality allows the ability to visualize the predicted placement of the unknown isolate in a larger phylogenetic context. The predicted shared ANI with the unknown isolate is displayed as a radial bar chart around the perimeter of the tree (fig. 3, external ring).
3. For the currently displayed tree, the abundance of represented species, phylogroups, and pathovars are to the left of the tree. (fig. 3)
4. A summary of the virulence factors found among the most closely related of the unknown isolate genomes given the selected ANI threshold is available as a second tab in the results section. (fig. 4)

**Figure 3.**
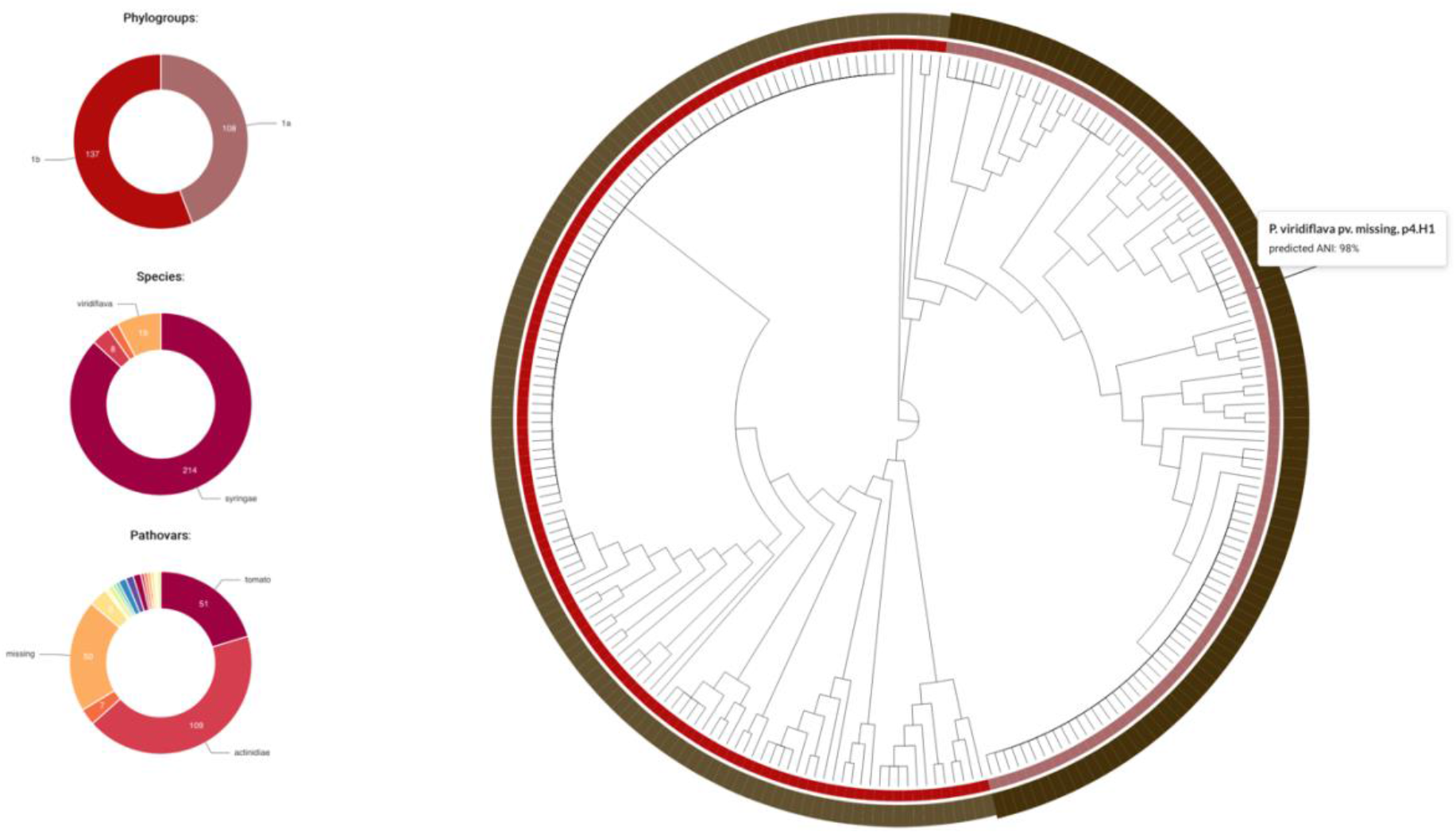
screenshot of classification results provided by Syringae.org. The tree is rooted at the most recent common ancestor of all genomes within the LIN cluster the unknown isolate has been placed into. The exterior ring shows the predicted ANI similarity between the unknown isolate and each reference genome. The interior ring shows phylogroup assigned to each reference genome. Relative abundance of phylogroups, species, and pathovars found within the unknown isolates predicted LIN cluster are shown to the left of the tree.

**Figure 4.**
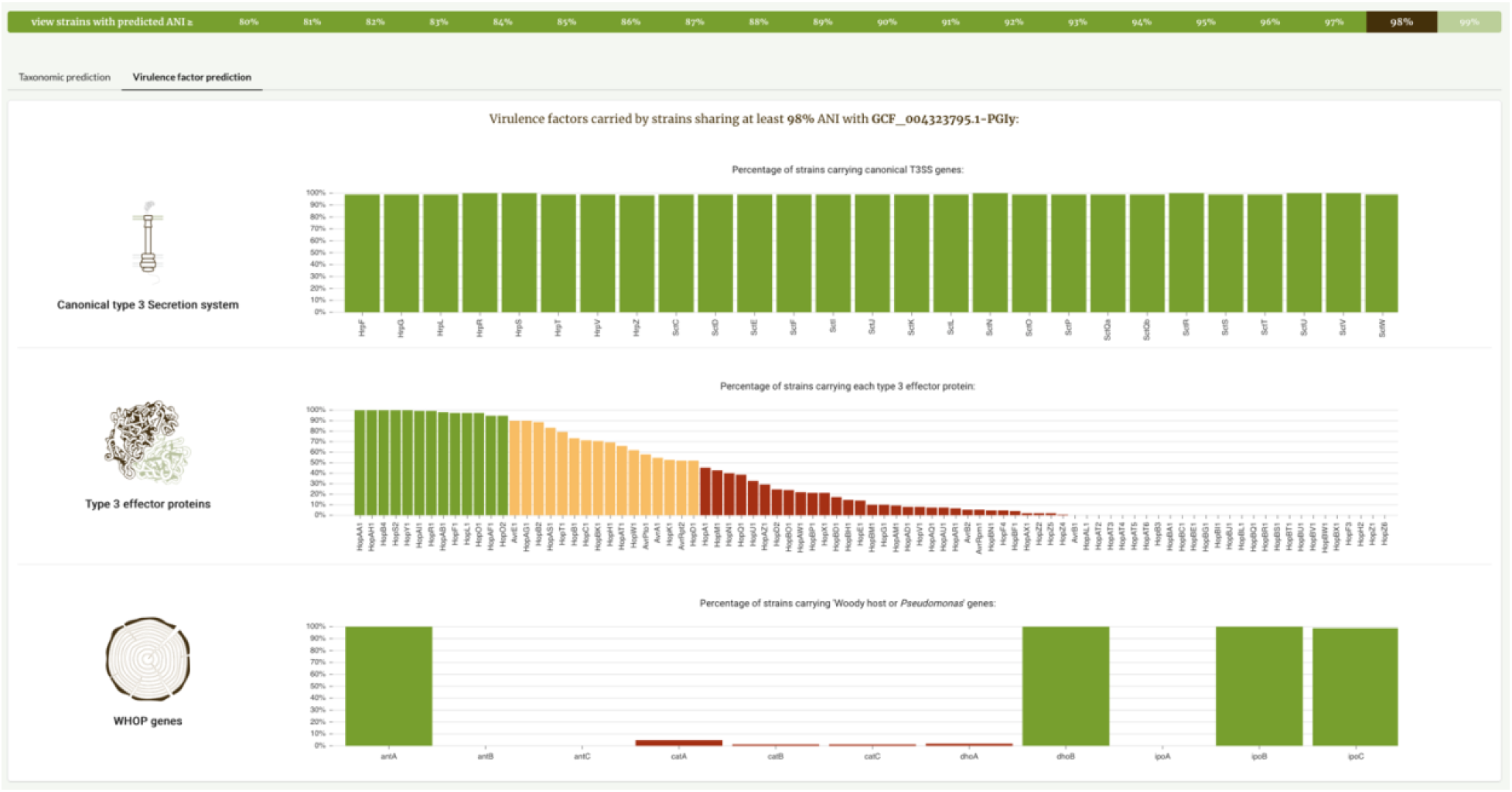
screenshot of virulence factor prediction provided by Syringae.org. The proportion of genomes within an unknown isolate’s predicted LIN cluster that carry canonical type III secretion system genes, type III effectors, and WHOP genes are displayed. Green, yellow, and red bars denote virulence factors found in >90%, >50%, and <50% of related genomes, respectively.

To quickly learn more about any genomes predicted to be similar to the unknown isolate, users can utilize the *Explore* and *Search* functionalities.

#### Explore

An interactive visualization tool for exploring the phylogenetic and genetic diversity of PSSC. Users can filter the 2,161-genome phylogenetic tree Syringae uses for visualizing classification data by taxa (phylogroup, species, and pathovar), and annotate up to six features, including multiple taxa and presence/absence of up to 3 NCBI-annotated genes (fig. 5)

**Figure 5.**
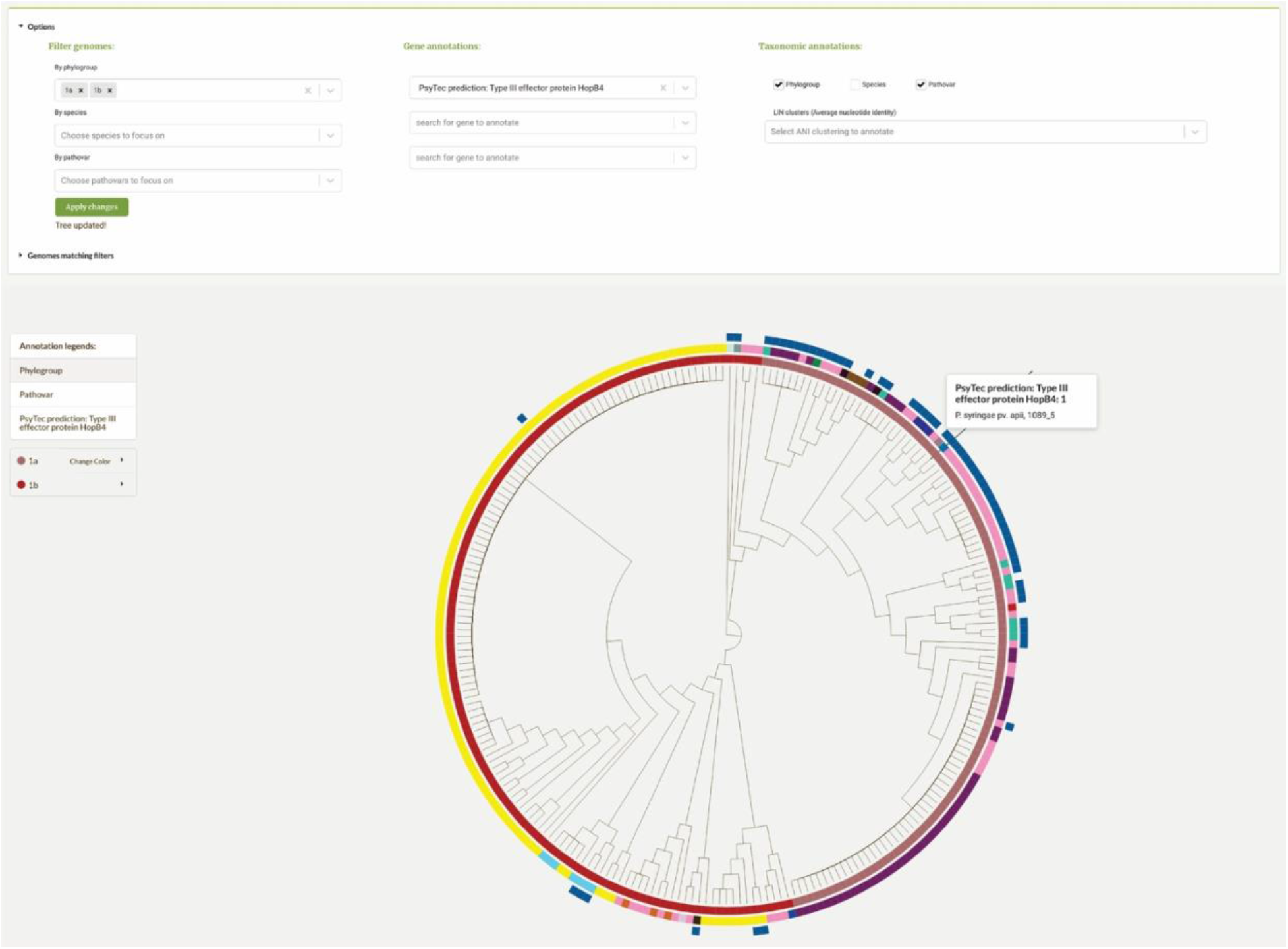
Screenshot of the Explore functionality on Syringae.org. The form at top allows for filtering of genomes to display by taxa, and annotation by up to 3 simultaneous genes and 4 taxa. Annotation rings surrounding the phylogenetic tree are based on user-selected annotations; in this case showing, from innermost ring outward, phylogroups, pathovars, and absence/presence of effector protein subfamily HopB4.

#### Search

A search tool for quickly finding metadata for any genome in SYRINGAE’s database. Search results include a list of the closest PSSC relatives in the dataset, as measured by fastANI (fig. 6). Users can also search for any NCBI-annotated gene names or virulence factors annotated by us for Syringae. Search results include a list of all genomes in our database carrying the gene of interest, as well as each unique protein accession number associated with the gene name found within the species complex.

**Figure 6.**
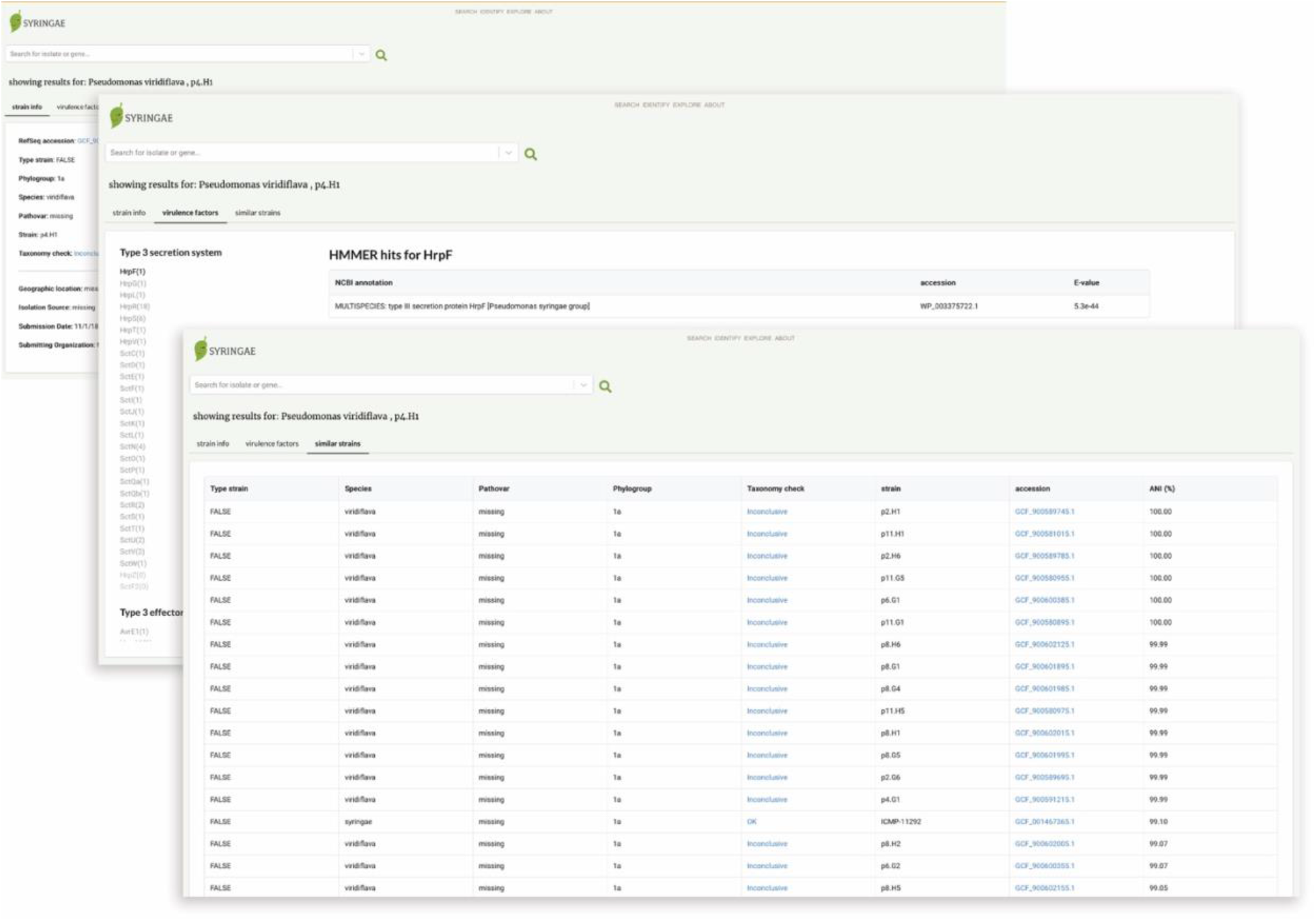
screenshot of search results for a PSSC genome displaying from top to bottom: metadata associated with the genome, predicted virulence factors, and most closely related genomes.

